# Changes in clock gene expression in esophagus in rat reflux esophagitis

**DOI:** 10.1101/489195

**Authors:** Atsushi Hashimoto, Risa Uemura, Akinari Sawada, Yuji Nadatani, Koji Otani, Shuhei Hosomi, Yasuaki Nagami, Fumio Tanaka, Noriko Kamata, Koichi Taira, Hirokazu Yamagami, Tetsuya Tanigawa, Toshio Watanabe, Yasuhiro Fujiwara

**Affiliations:** Department of Gastroenterology, Osaka City University Graduate School of Medicine, Osaka, Japan.

## Abstract

**Background and Aims:** Gastroesophageal reflux disease (GERD) is strongly associated with sleep disturbances. Clock genes harmonize circadian rhythms by their periodic expression and regulate several physiological functions. However, the association between clock genes and GERD is still unknown. We investigated whether reflux esophagitis affects circadian variability of clock genes in the esophagus and other organs using a rat reflux esophagitis model.

**Methods:** Reflux esophagitis was induced in 7-week-old male Wistar rats. Sham-operated rats were used as controls. Rats were sacrificed at 09:00 (light period) and 21:00 (dark period) 3 days (acute phase) and 21 days (chronic phase) after induction of esophagitis. The expression levels of clock gene mRNAs such as *Per1*, *Per2*, *Per3*, *Cryl*, *Cry2*, *Arntl*, and *Clock* in the esophagus were investigated by qPCR. *Arntl* expression was examined in stomach, small intestine, colon, and liver tissues. Serum melatonin and IL-6 levels were measured by ELISA.

**Results:** Histological examination of reflux esophagitis mainly revealed epithelial defects with marked inflammatory cell infiltration in the acute phase, and mucosal thickening with basal cell hyperplasia in the chronic phase. Circadian variability of clock genes, except *Cry1,* was present in the normal esophagus, and was completely disrupted in reflux esophagitis during the acute phase. The circadian variability of *Per2*, *Per3*, and *Arntl* returned to normal, but disruption of *Per1*, *Cry2*, and *Clock* was present in the chronic phase. Disruption of circadian variability of *Arntl* was observed in the esophagus, as well as in the stomach, small intestine, and liver tissues in reflux esophagitis during the acute phase. There were no significant differences in serum melatonin and IL-6 levels between control and reflux esophagitis animals in both acute and chronic phases.

**Conclusion:** Disruption to circadian variability of clock genes may play a role in the pathogenesis of GERD.

## Introduction

Gastroesophageal reflux disease (GERD) is a common gastrointestinal (GI) disorder. GERD is caused by the abnormal reflux of the gastric contents, with reflux symptoms such as heartburn [1]. Several clinical studies reported that GERD is strongly associated with sleep disturbances [2-5]. Our previous animal study confirmed that rat reflux esophagitis is associated with an increase in wakefulness, accompanied by a reduction in non-rapid eye movement (NREM) sleep during light periods, an increase in sleep fragmentation, and more frequent stage transitions [6].

Clock genes induce circadian rhythm by their periodic expression and regulate several physiological functions [7-9]. The mammalian circadian rhythm is generated by a molecular circadian-clock system, involving genes including periods (*Per1*, *Per2*, and *Per3*), cryptochromes (*Cry1* and *Cry2*), *Arntl* (Aryl hydrocarbon receptor nuclear translocator like 1, aka *Bmal1*), and *Clock* (circadian locomotor output cycles kaput) [7-11]. This clock system involves a feedback system that depends on two core clock proteins, CLOCK and BMAL1. These proteins heterodimerize in cytoplasm, forming a complex that activates their target genes, Periods and Cryptochromes, which form a repressor complex that interacts with CLOCK-BMAL1, resulting in inhibition of their own transcription [12, 13]. Several studies showed that the circadian rhythms of clock genes are associated with sleep conditions [11, 14], and are present in the whole body, including the central circadian pacemaker, the suprachiasmatic nucleus, and many peripheral tissues, such as heart, liver and gastrointestinal (GI) tract [15, 16].

Although the circadian-clock system is susceptible to inflammation [17], detailed associations between clock genes and GERD have not been elucidated. We investigated whether reflux esophagitis affects circadian variability of clock genes using a rat reflux esophagitis model.

## Methods

### Animals and induction of esophagitis

The experimental protocols involved in this study were approved by the Institutional Animal Care and Use Committee of the Osaka City University Graduate School of Medicine. Specific pathogen free male Wistar rats (Japan SLC, Hamamatsu, Japan) weighing approximately 180 g at the start of the experiment were used. They were housed at a constant temperature (22 ± 2°C) in cages with an automatically controlled 12 h light/dark cycle (light on at 08:00 and light off at 20:00) and they had free access to food and water. Reflux esophagitis was induced by the methods described by Omura and colleagues [18]. In brief, the duodenum near the pyloric ring was covered with a 2 mm-wide piece of 18 Fr Nelaton catheter (Terumo Co, Tokyo, Japan), and the transitional region between the forestomach and the glandular portion was ligated to enhance reflux of gastric contents into the esophagus. Solid food was withdrawn for two days after induction of esophagitis, but rats were allowed drinking water. Sham-operated rats were used as controls. Rats were sacrificed humanely at 09:00 (light period) and 21:00 (dark period) 3 days (acute phase) and 21 days (chronic phase) after induction of esophagitis. Esophageal lesions were examined macroscopically and histologically. Esophagus, stomach, small intestine, colon, and liver tissues were excised and immediately stored in RNA later solution (Applied Biosystems, Foster, CA, USA) until real time quantitative polymerase chain reaction (qPCR) was performed. For histological analysis, samples were gently rinsed with saline and fixed in 10% buffered formalin. Samples were embedded in paraffin and cut to 4 μm thick sections. Hematoxylin and eosin staining was performed for standard morphological analysis. Blood samples from rats were collected by puncture during terminal anesthesia and centrifuged at 1500 × g for 10 min at 4 °C. Supernatants were transferred to fresh tubes and stored at −80°C until analysis.

### RNA isolation and qPCR

Total RNA was extracted and purified from frozen gastrointestinal tissues using the ISOGEN kit (Nippon Gene Co., Ltd., Tokyo, Japan). Complementary DNA was produced using the High Capacity RNA-to-cDNA Kit (Thermo Fisher Scientific Inc., Waltham, MA, USA). qPCR analyses were performed using an Applied Biosystems 7500 Fast Real-Time PCR system and software (Thermo Fisher Scientific Inc.). Thermal cycling conditions were 45 cycles of amplification at 95°C for 15 seconds and 60°C for one minute. PCR primers and TaqMan probes for *Per1*, *Per2*, *Per3*, *Cry1*, *Cry2*, *Arntl*, and *Clock* were used. The primer sequences were shown in Table 1. Expression levels were normalized to those of TaqMan glyceraldehyde-3-phosphate dehydrogenase (*GAPDH*; Thermo Fisher Scientific Inc.) mRNA levels.

**Table 1.**
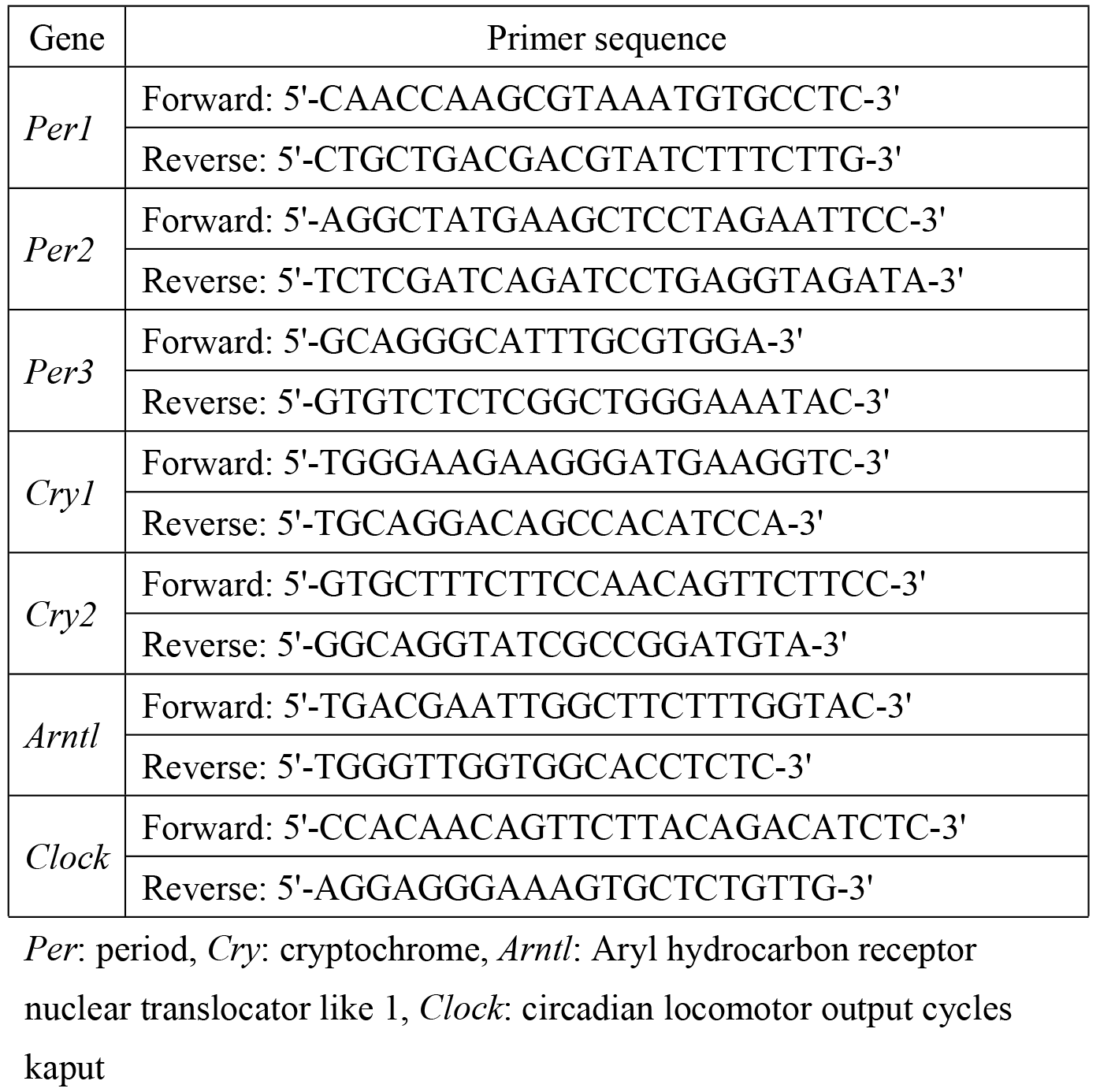
Sequences of primers for q-PCR.

### Enzyme-linked immunosorbent assays (ELISA) for serum melatonin and IL-6

The amounts of melatonin and IL-6 in serum proteins were measured using the Melatonin ELISA Kit (Enzo Life Sciences, Inc., New York, NY, USA) and Interleukin 6 ELISA Kit (Cusabio Technology, Houston, TX, USA), respectively, following the manufacturers’ instructions.

### Statistical analysis

Data were analyzed by Kruskal-Wallis test (Steel Dwass post-hoc comparison) for statistical comparison between each group. The level of statistical significance was set to p < 0.05. All statistical analyses were performed using the statistical software EZR (Easy R), which is based on R and R commander [19].

## Results

### Macroscopic and histological assessment

The macroscopic and histological appearance of the normal esophagus and reflux esophagitis in the rat model are shown in Fig 1. Normal esophagus had a thin epithelial layer, with squamous cells and few inflammatory cells in the submucosal layer (Fig 1-A, D). In contrast, there were multiple erosions and edematous mucosa on day 3 (Fig 1-B). Histological defects of the epithelial layer, with marked infiltration of inflammatory cells in the lamina propria, submucosal layer, and bases of ulcers were observed (Fig 1-E). There were whitish lesions in the middle esophagus with edematous surfaces on day 21 (Fig 1-C). Mucosal thickening, with elongation of the lamina propria papillae and basal cell hyperplasia and infiltration of inflammatory cells, was observed (Fig 1-F).

**Fig 1.**
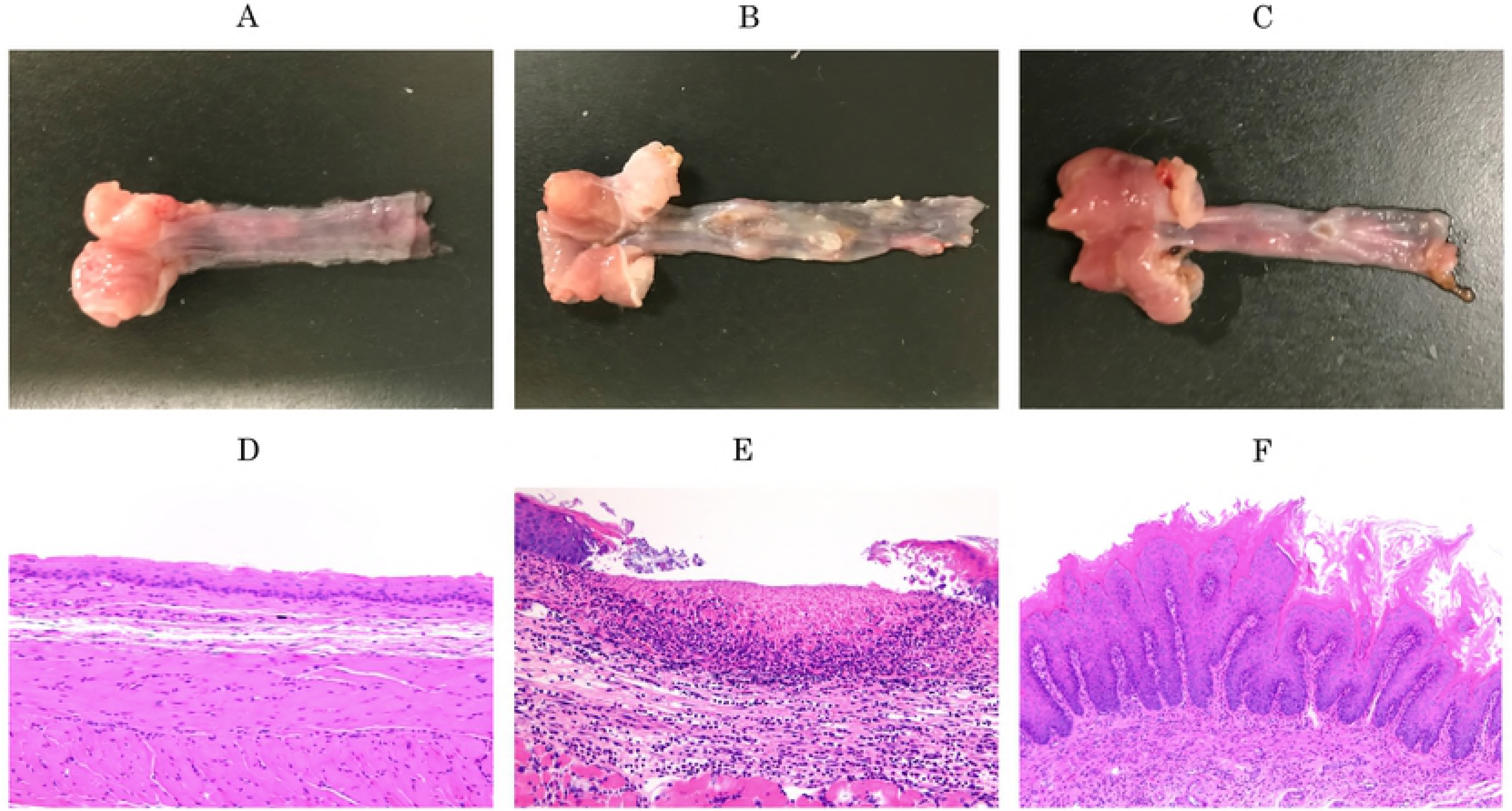
Macroscopic (A, B, C) and histological (E, F, G) appearances of normal esophagus and reflux esophagitis in the rat model.

### Circadian variability of clock gene expression in rat esophagus

Fig 2 presents the mRNA expression levels of *Per1*, *Per2*, *Per3*, *Cry2*, *Arntl*, and *Clock* in rat esophagus. In controls, mRNA expression levels of *Per1*, *Per2*, *Per3*, and *Cry2* showed circadian variability, which was lower at 09:00 and higher at 21:00 during the acute and chronic phases, respectively (Fig 2-A, B, C, D). In contrast, mRNA of *Arntl* and *Clock* exhibited circadian variability which was higher at 09:00 and lower at 21:00 in the acute and chronic phases, respectively (Fig 2-E, F). The pattern of circadian variability of *Per1*, *Per2*, *Per3*, and *Cry2* expression was opposite to that of *Arntl* and *Clock*. In reflux esophagitis, there were no difference in *Per1*, *Per2*, *Per3*, *Cry2*, *Arntl,* and *Clock* expression between light and dark periods during the acute phase, and in *Per1*, *Cry2*, and *Clock* expression in the chronic phase (Fig 2). The mRNA expression levels of *Cry1* had no circadian variability in both control and reflux esophagitis during acute and chronic phases (data not shown).

**Fig 2.**
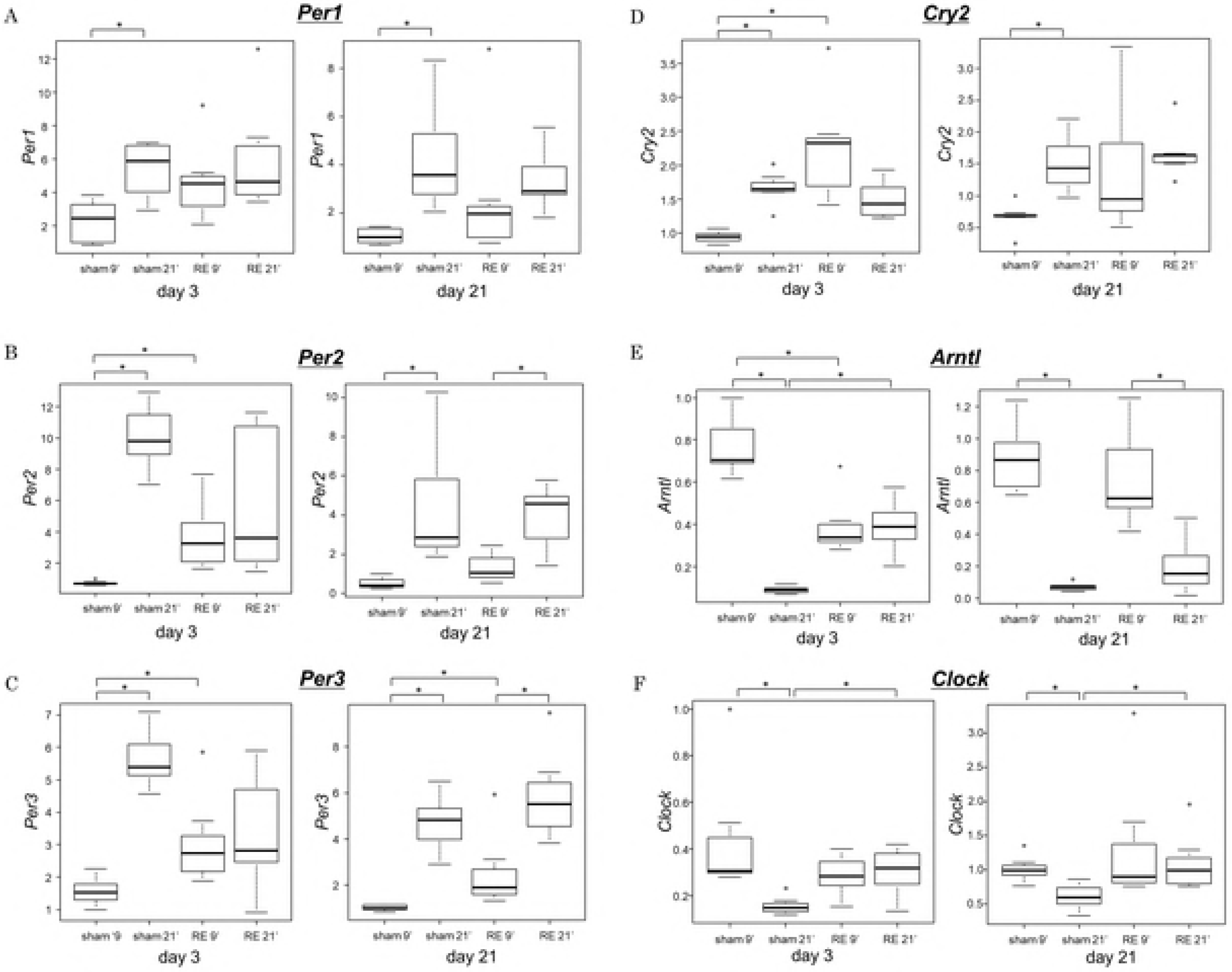
The mRNA expression levels of *Per1*, *Per2*, *Per3*, *CRY2*, *Arntl* and *Clock* in (1) sham-operated rats sacrificed at 09:00, (2) sham-operated rats sacrificed at 21:00, (3) reflux esophagitis (RE) rats sacrificed at 09:00 and (4) reflux esophagitis rats sacrificed at 21:00 during the acute phase (day 3) and chronic phase (day 21), respectively. *p < 0.05.

The circadian variability of clock genes in the present study is summarized in Table 2. The normal esophagus showed circadian variability in *Per1*, *Per2*, *Per3*, *Cry2*, *Arntl*, and *Clock* genes, but reflux esophagitis disrupted circadian variability of each gene during the acute phase. The disrupted circadian variability was improved in *Per2*, *Per3*, and *Arntl,* but not in *Per1*, *Cry2*, and *Clock,* in the chronic phase.

**Table 2.** Circadian variability in clock genes of rat esophagus.

|  | Acute phase |  | Chronic phase |  |
| --- | --- | --- | --- | --- |
|  | Control | RE | Control | RE |
| *Per1* | (+) | (-) | (+) | (-) |
| *Per2* | (+) | (-) | (+) | (+) |
| *Per3* | (+) | (-) | (+) | (+) |
| *Cry1* | (-) | (-) | (-) | (-) |
| *Cry2* | (+) | (-) | (+) | (-) |
| *Arntl* | (+) | (-) | (+) | (+) |
| *Clock* | (+) | (-) | (+) | (-) |

RE: reflux esophagitis, *Per*: period, *Cry*: cryptochrome, *Arntl*: Aryl hydrocarbon receptor nuclear translocator like 1, *Clock*: circadian locomotor output cycles kaput

### Circadian variability in various organs

Fig 3 shows mRNA expression levels of *Arntl* in the esophagus, stomach, small intestine, colon, and liver tissues. Control groups showed circadian variability of *Arntl* genes in the esophagus as well as in stomach, small intestine, colon, and liver, whereas circadian variability was disrupted in organs other than colon in reflux esophagitis during the acute phase.

**Fig 3.**
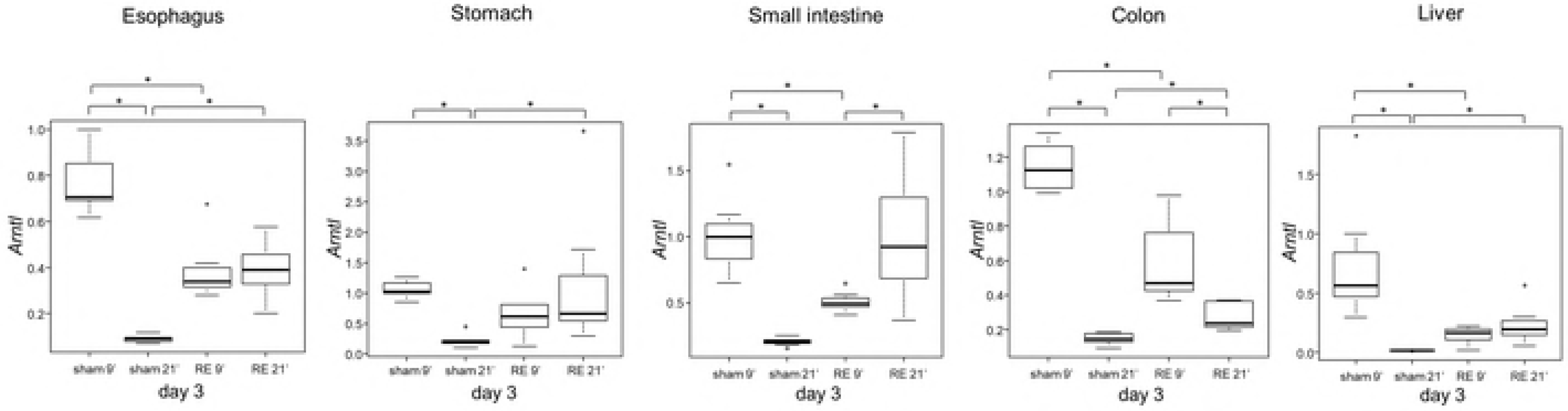
The mRNA expression levels of *Arntl* in the esophagus, stomach, small intestine, colon, and liver tissues in controls and reflux esophagitis (RE). *p<0.05.

### Serum melatonin and IL-6 levels

Fig 4 shows serum melatonin and IL-6 levels in control and reflux esophagitis in the acute and chronic phases. There were no significant differences in serum levels of melatonin between 09:00 and 21:00, suggesting that serum melatonin had no circadian variability. Furthermore, there were no differences in serum melatonin levels between control and reflux esophagitis (Fig 4-A). The serum IL-6 level was significantly higher at 09:00, compared to 21:00, but only in controls in the chronic phase. However, there were no differences in serum IL-6 levels between control and reflux esophagitis (Fig 4-B).

**Fig 4.**
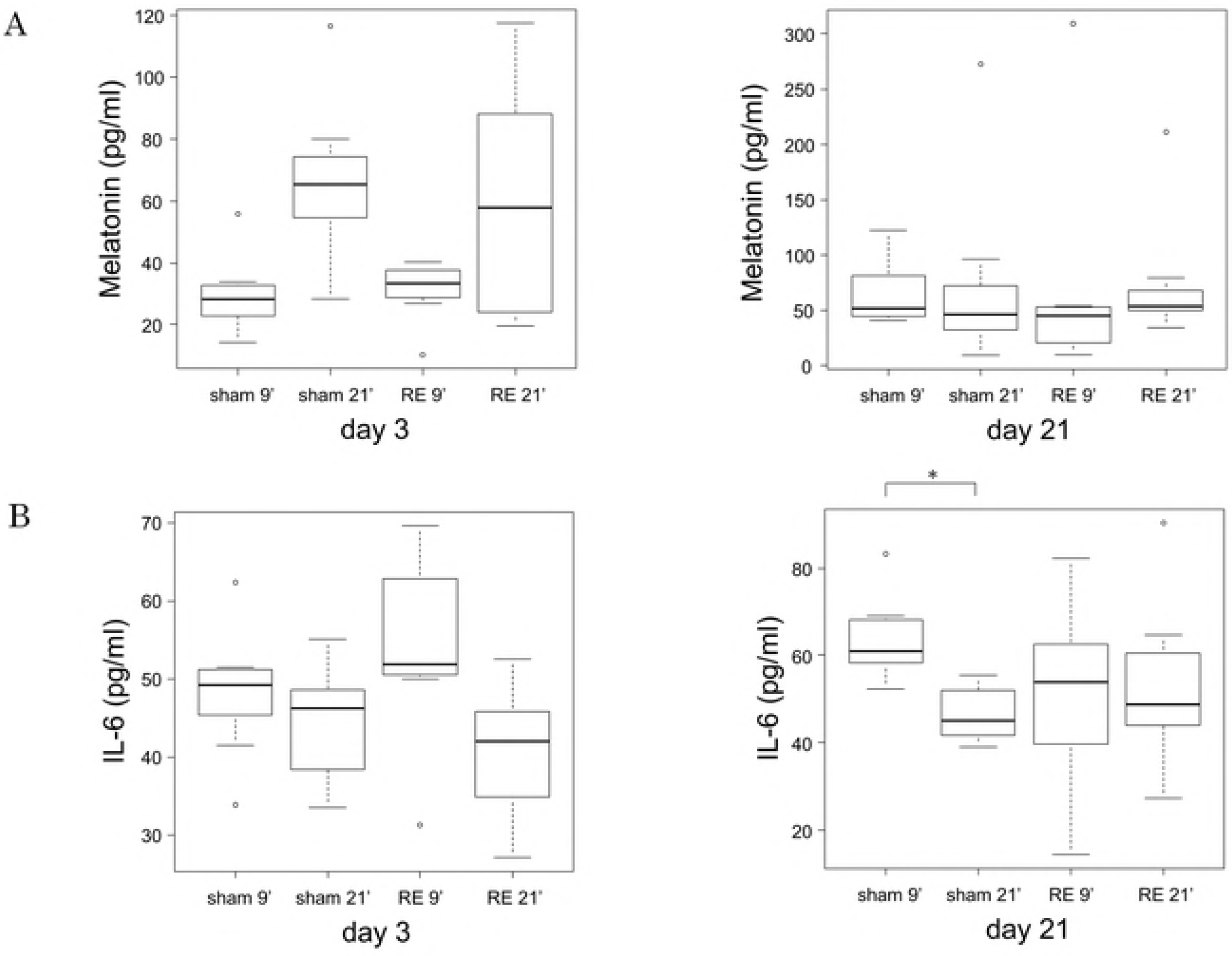
Serum melatonin and IL-6 levels in control groups and in reflux esophagitis groups during the acute and chronic phase. *p < 0.05.

## Discussion

The present study demonstrated that circadian variability of clock genes, except *Cry1,* was present in the normal esophagus, and that these circadian variabilities were completely disrupted in reflux esophagitis during the acute phase. Furthermore, disruption of circadian variability of *Per1*, *Cry2* and *Clock* was still present, but other genes, such as *Per2*, *Per3* and *Arntl,* returned to circadian variability like normal esophagus. The circadian variability of *Arntl* was disrupted in the esophagus as well as stomach, small intestine, and liver tissues in reflux esophagitis in the acute phase. These results suggested that changes in clock gene expression might play a role in the pathogenesis of GERD and be related to expression of clock genes elsewhere in the GI tract, and in liver.

This study is the first detailed analysis regarding clock genes in rat esophagus. The present study showed that expression of *Per1*, *Per2*, *Per3*, and *Cry2* mRNA at 09:00 (light period) was low, and high at 21:00 (dark period), while expression of *Arntl* and *Clock* was high at 09:00 and low at 21:00 in normal esophagus, suggesting the presence of circadian rhythms in clock genes in the esophagus. The expression of Periods and Cryptochromes indicate an opposite pattern to the expression of *Arntl* and *Clock* due to a negative feedback relationship in the clock cycle [12, 13]. Several studies reported that other organs, such as hypothalamic nuclei, adrenal gland, liver, heart, kidney and spleen revealed similar patterns of clock gene expression [20-22].

Circadian variability of most clock genes was disrupted in the reflux esophagitis in the acute phase, whereas several clock genes, such as *Per2*, *Per3*, and *Arntl*, showed circadian variability, similar to normal esophagus in the chronic phase. These results suggest that the circadian rhythm of clock genes is susceptible to acute inflammation, compared to chronic inflammation. Although several studies reported associations between clock genes and chronic inflammatory diseases, including rheumatoid arthritis, allergies, and inflammatory bowel disease (IBD) [23-25], there are no reports concerning associations between clock genes and acute inflammation.

Yang et al. investigated clock genes in healthy controls and patients with GERD [26]. The authors showed that the rhythmic patterns and levels of *Per1*, *Per2*, *Bmal1*, and *Cry2* expression in the esophagus were disrupted in patients with GERD. Although *Per1* and *Per2* expression in the esophagus of both controls and GERD patients showed similar patterns of circadian variability, expression of *Bmal1* and *Cry2* in GERD revealed a reverse pattern of control. We identified disruption of circadian variability of clock genes, such as *Per1*, *Cry2*, and *Clock*, in reflux esophagitis during the chronic phase, but an inverse pattern of such genes was not observed, and *Per2*, *Per3* and *Arntl* exhibited circadian variability, comparable to normal esophagus. Discrepancies between the present study and Yang’s might be due to study subjects (human versus rats), and to the severity of reflux esophagitis.

Several studies reported associations between clock genes and chronic inflammation [17, 27-29]. In the gastrointestinal field, several studies showed that disruption of the circadian variability of clock genes can cause immune activation, and release inflammatory cytokines, resulting in sleep disturbances in IBD [27, 30, 31]. Since reflux esophagitis induces increases in levels of cytokines such as IL-1β, IL-6, and IL-8 in the esophageal mucosa [32], disruption of circadian variability of clock genes might affect expression of these cytokines in GERD.

Circadian variability of *Arntl* was observed in the esophagus, stomach, small intestine, colon, and liver in controls, and was disrupted in the esophagus as well as stomach, small intestine, and liver in reflux esophagitis. We predicted two mechanisms by which reflux esophagitis disrupted circadian variability of clock genes in other organs. First, sleep disturbance caused by reflux esophagitis affects core clock genes in the suprachiasmatic nucleus, and the central clock system synchronize peripheral tissues. Second, reflux esophagitis primarily disrupts circadian variabilities of clock genes in the esophagus, and affects the central clock system, resulting in synchronization in other organs. Since serum melatonin and IL-6 levels showed no differences between control and reflux esophagitis groups, melatonin and IL-6 are unlikely to synchronize circadian variability. Therefore, other hormonal factors or the autonomic nervous system might play a role in synchronization.

Mechanisms of how clock genes affect GERD pathogenesis should be discussed. Our study showed that *Per2*, *Per3*, and *Arntl* expression returned to normal circadian variability, but disruption of circadian variability of *Per1*, *Cry2*, and *Clock* was present in reflux esophagitis during the chronic phase. Although the exact role of each clock gene in GERD pathogenesis is unknown, it is interesting that the pattern of clock gene expression changed during transition of the acute phase to the chronic phase.

Histologically, there was marked infiltration of inflammatory cells in the acute phase and mucosal thickening with basal cell hyperplasia during the chronic phase, suggesting that the chronic phase reveals an epithelial repair phase. Since an association between wound healing in the skin and patterns of clock gene expression has been reported [33], clock genes might affect epithelial proliferation through cell cycle control in reflux esophagitis.

In conclusion, circadian variability of several clock genes was disrupted in reflux esophagitis, especially during the acute phase, and reflux esophagitis affected expression of clock genes elsewhere in the GI tract and liver. These findings suggest that changes in clock gene expression might play a role in the pathogenesis of GERD.

## Acknowledgments

We thank Emi Yoshioka for her technical assistance.

Author contributions
Study concept and design: A.H. and Y.F.; acquisition of data: A.H., R.U., A.S. and Y.F.; analysis and interpretation of data: A.H. and Y.F.; drafting of manuscript: A.H.; critical revision of the manuscript for important intellectual content and approval of the final version: A.H., R.U., A.S., Y.Nadatani., K.O., K.T., S.H., Y.Nagami., F.T., N.K., H.Y., T.T., T.W., and Y.F.

## Notes

**Potential competing interests**: None

